# Leveraging distant relatedness to quantify human mutation and gene conversion rates

**DOI:** 10.1101/020776

**Authors:** Pier Francesco Palamara, Laurent Francioli, Giulio Genovese, Peter Wilton, Alexander Gusev, Hilary Finucane, Sriram Sankararaman, The Genome of the Netherlands Consortium, Shamil Sunyaev, Paul I.W. de Bakker, John Wakeley, Itsik Pe’er, Alkes L. Price

## Abstract

The rate at which human genomes mutate is a central biological parameter that has many implications for our ability to understand demographic and evolutionary phenomena. We present a method for inferring mutation and gene conversion rates using the number of sequence differences observed in identical-by-descent (IBD) segments together with a reconstructed model of recent population size history. This approach is robust to, and can quantify, the presence of substantial genotyping error, as validated in coalescent simulations. We applied the method to 498 trio-phased Dutch individuals from the Genome of the Netherlands (GoNL) project, sequenced at an average depth of 13x. We infer a point mutation rate of 1.66 ± 0.04 × 10^−8^ per base per generation, and a rate of 1.26 ± 0.06 × 10^−9^ for < 20 bp indels. Our estimated average genome-wide mutation rate is higher than most pedigree-based estimates reported thus far, but lower than estimates obtained using substitution rates across primates. By quantifying how estimates vary as a function of allele frequency, we infer the probability that a site is involved in non-crossover gene conversion as 5.99 ± 0.69 × 10^−6^, consistent with recent reports. We find that recombination does not have observable mutagenic effects after gene conversion is accounted for, and that local gene conversion rates reflect recombination rates. We detect a strong enrichment for recent deleterious variation among mismatching variants found within IBD regions, and observe summary statistics of local IBD sharing to closely match previously proposed metrics of background selection, but find no significant effects of selection on our estimates of mutation rate. We detect no evidence for strong variation of mutation rates in a number of genomic annotations obtained from several recent studies.

## Introduction

Germline mutations represent a fundamental evolutionary force that shapes phenotypic variation and has a profound impact on heritable diversity. Precise estimation of mutation rates has several applications, including the interpretation of mutations implicated in diseases [1, 2, 3, 4], studies of natural selection [5, 6, 7], the timing of demographic events inferred using genetic analysis [8, 9, 10, 11], and the study of several aspects of human mutagenesis [12]. High throughput sequencing technologies have recently enabled the quantification of germline mutation rates, but the estimates obtained using these methods are inconsistent with previous studies. The source of these inconsistencies, whether biological or due to methodological biases, is at the center of recent debate [10], and new methods are required to gain additional insight into germline mutation rates.

Several aspects of methods for inferring mutation rates were recently reviewed in [13, 14]. Approaches based on sequence divergence among primates [15, 16] (phylogenetic methods) estimate mutation rates that range between 2.0 − 2.5 × 10^−8^. These methods depend on a number of population-genetic assumptions, and further uncertainty is contributed by the need to map inferred per-year mutation rates to a per-generation scale. Pedigree-based estimates of mutation rates ranging between 0.9 − 1.3 × 10^−8^ [17, 18, 19, 3, 20, 21], on the other hand, are based on direct observation of mutation events, but issues related to sequencing error complicate the inference, requiring the use of stringent genotyping filters. Several statistical inconsistencies, such as underdispersion of the reported estimates, have been outlined in [13]. This motivates the development of new methods for estimating mutation rates. Two methods based on the pairwise sequentially Markovian coalescent (PSMC [9]) recently estimated slightly higher mutation rates than pedigree-based studies ([22, 23]).

In this work, we propose a new method for estimating mutation rates using mutations occurring within identical-by-descent haplotype blocks (IBD [24, 25, 26, 27, 28, 29, 30, 31, 32]) transmitted through recent common ancestors that lived in the past ∼100 generations (∼3. 000 years) before present. Our approach is robust to, and can quantify, the presence of substantial amounts of genotyping error in the analyzed sequences. By quantifying how estimates vary as a function of allele frequency, we correct our estimate for additional biases created by the occurrence of gene conversion events on the ancestral lineages leading to shared common ancestors. We apply this methodology to analyze 250 trio families from the Netherlands sequenced at an average of ∼13×, obtaining a genome-wide average point mutation rate estimate of 1.66 ± 0.04 × 10^−8^ per base, per generation. We further apply our methodology to infer the rate of < 20 bp indels, which we estimated to be 1.26 ± 0.06 × 10^−9^. We analyze the relationship between recombination rates and mutation rates [33], and find that after accounting for the occurrence of gene conversion events, no significant association is detected. In addition to estimating the rate of mutation events, we derived an estimate for the rate at which a genomic locus is involved in a non-crossover gene conversion event of 5.99 ± 0.69 × 10^−6^. We find that mismatching variants within IBD regions - representing mutations of recent origin - are strongly enriched for deleterious variation. We show that the length of IBD sharing along the genome closely reflects widely adopted summary statistics of background selection [5]. Background selection, however, is observed not to have significant effects on our estimates. Finally, we explore enrichment or depletion of several specific genomic annotations, and find no evidence for substantial differences compared to the average genome-wide mutation rate.

## Materials and Methods

### Overview of methods

Pairs of purportedly unrelated individuals from a population often share long stretches of chromosomal regions inherited identical-by-descent (IBD) from recent common ancestors that lived in the past tens to few hundreds of generations. These IBD segments can be detected using several available methods [34, 25, 35, 36], and reflect genetic relationships that are typically not known to the affected individuals, but are found to be ubiquitous even in outbred populations [37, 30]. IBD segments are defined in our work as contiguous chromosomal regions for which two sampled chromosomes share the same most recent common ancestor (MRCA). In this work, we are interested in IBD segments co-inherited from ancestors that lived within ∼100 generations before present, which can be reliably detected in trio-phased real data. Occasional mutations segregating along the lineages connecting a pair of IBD haplotypes to their MRCA will create mismatched sites on the shared haplotypes that can be used to infer the rate at which new germline mutations appear. If the exact number of generations separating the IBD segments (via their MRCA) is known, one may infer the mutation rate by dividing the number of observed sequence mismatches by the number of generations and the physical length, for all segments. A special case of this approach is used in trio-based analyses, where transmitted parental haplotypes and IBD offspring haplotypes are separated by a single generation.

We briefly describe solutions to three challenges. First, to estimate the number of generations separating two IBD segments (twice the time to most recent common ancestor; tMRCA), we use a recently developed method [29] that relies on the spectrum of observed IBD segment lengths to infer demographic history, which is then used to obtain a posterior mean estimate of the average tMRCA for pools of IBD segments of different lengths, as detailed in the real data description and the Appendix. Second, to deal with the presence of genotyping errors, rather than relying on stringent filtering criteria (as in trio-based analyses [17, 18, 19, 3, 20, 21]), we regress the observed sequence mismatches for several IBD length thresholds on the estimated tMRCA; the slope of this regression reflects the rate at which new mutations accumulate per generation time unit, while the genotyping error rate is captured by the intercept. We refer to this procedure as tMRCA regression (illustrated in Figure 1). Finally, we correct for the occurrence of noncrossover gene conversion events along the lineages leading to the MRCA, exploiting the relationship between an allele’s frequency and the probability that it is involved in a gene conversion event. To do this, we repeatedly perform tMRCA regression, only considering mismatching sites with frequency less than a specified maximum allele frequency (MaAF) value, and regress the obtained mutation rate estimates on the MaAF threshold. We show that the intercept of this regression provides a gene conversion-corrected mutation rate estimate. We refer to this procedure as MaAF-threshold regression (illustrated in Figure 2). This also allows us to estimate the rate at which a genomic locus is involved in a noncrossover gene conversion event, which is proportional to the difference between corrected and uncorrected estimates for mutation rates. We have released open-source software (IBDMUT) implementing these methods (see Web Resources).

**Figure 1:**
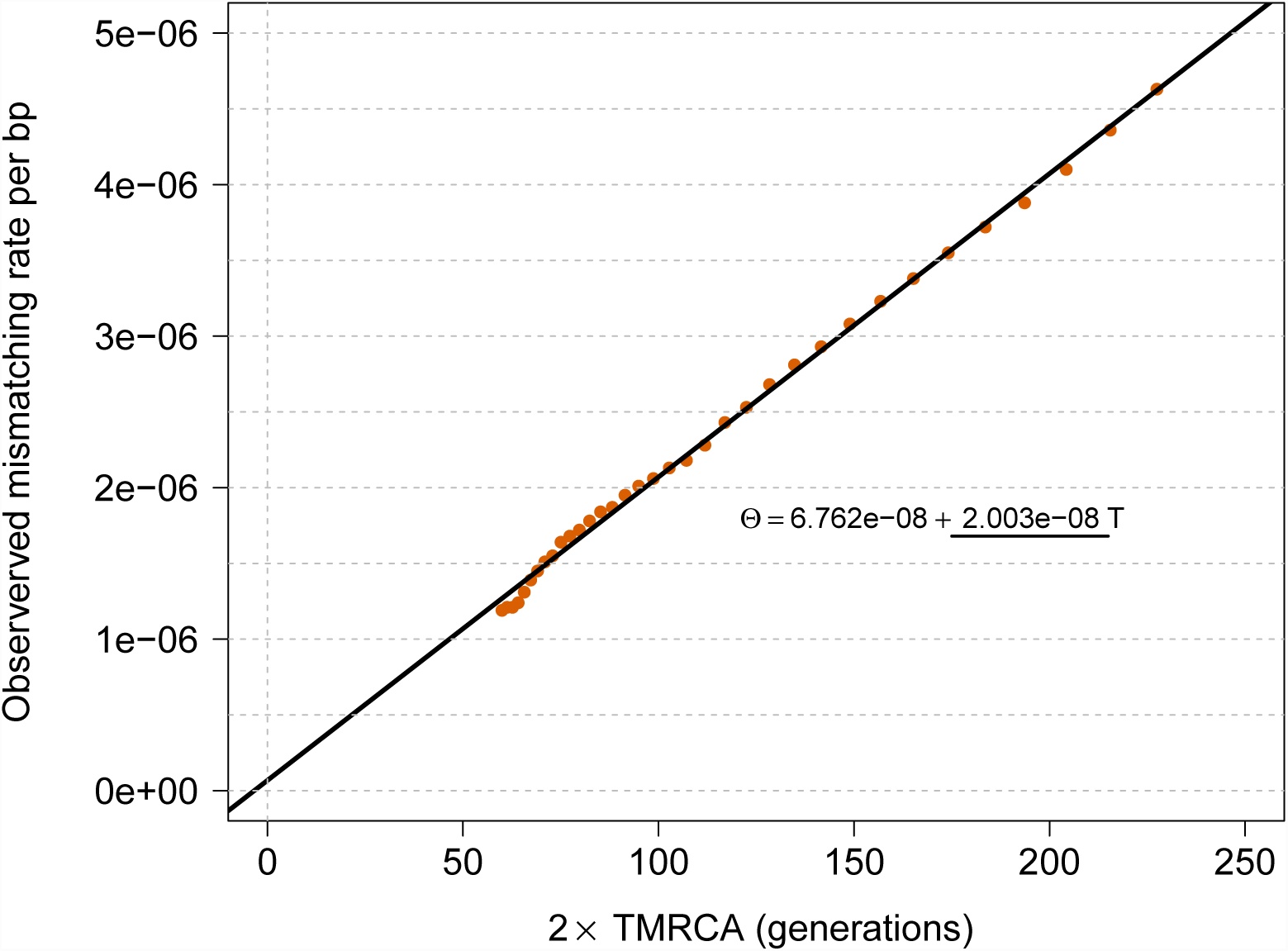
tMRCA regression. We simulated a chromosome of 50 cM for 250 diploid samples, using *μ* = 2 ×10^−8^ for the mutation rate and no genotyping error. We matched the allele frequency spectrum of the simulated samples to the spectrum found in real data for IBD detection with GERMLINE, and used the IBD detection parameters used in real data. The slope of this regression captures the simulated mutation rate; the intercept is proportional to genotyping error rate.

**Figure 2:**
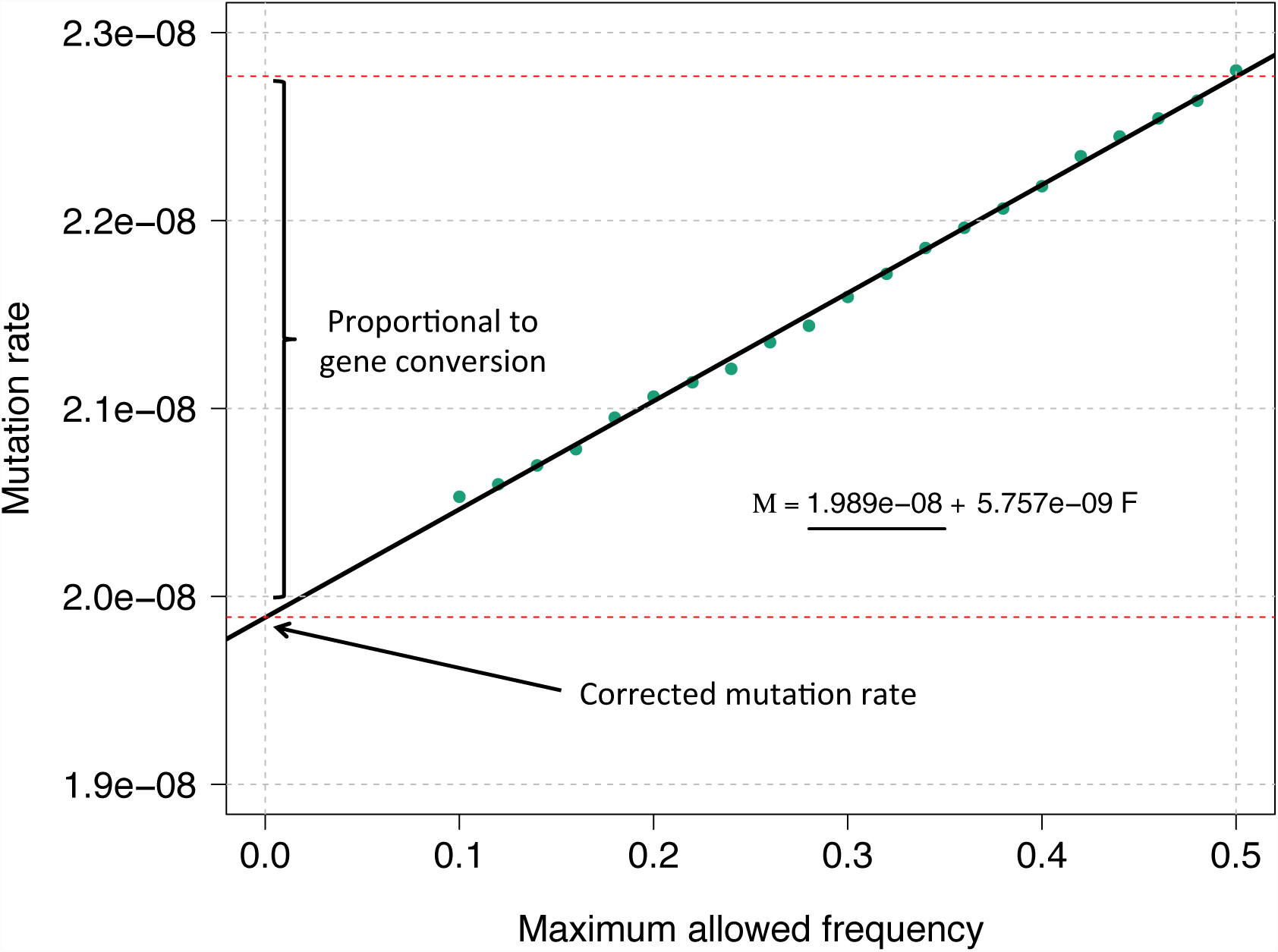
MaAF-threshold regression. We simulated 250 diploid samples as described in Figure 1, and a probability of 6 × 10^−6^ for a basepair to be involved in a non-crossover gene conversion event. We performed the MaAF-threshold regression to correct for the occurrence of gene conversion. The regression intercept is used to estimate the corrected mutation rate, while the difference between corrected and uncorrected mutation rates captures the effects of gene conversion, whose magnitude can be estimated using the observed population heterozygosity.

### IBD detection and demographic inference

Shared IBD segments can be detected in large samples of purportedly unrelated individuals using several available algorithms [34, 25, 35, 36]. These approaches typically rely on the similarity of haplotypes within pairs of individuals (identity-by-state), and probabilistic models that enable confidently detecting long (e.g. > 1 cM) shared IBD segments. The accuracy of IBD detection is substantially improved when trio-phased individuals are available, as is the case in the analyzed data set. For both simulations and real data analysis reported in this paper, we used the GERMLINE [25] IBD detection algorithm. Inferring demographic history for the past ∼100 generations can be achieved by analyzing the length distributions of shared IBD segments, which provide information on the distribution of recent coalescent events within a population [29, 38, 30]. Details of the IBD detection and demographic inference methods for the reported real data analysis are described in the “GoNL data set” section. In order to test the accuracy of the proposed methodology, we relied on “ground truth” IBD sharing in some simulations, i.e. IBD segments extracted from the sampled ancestral recombination graphs (ARG), and made use of the simulated demographic history to inform mutation and gene conversion rate inference in several synthetic scenarios.

### Estimating the mutation rate via tMRCA regression

The proposed methodology for the inference of mutation rates requires the availability of haploid genotype data, a list of IBD segments between pairs of haploid individuals that are longer than a specified Morgan length threshold, including start/end position, and a demographic model, which may be inferred from the spectrum of IBD shared segments as described in [29]. For each IBD segment *i*, we obtain an observed mismatch rate by counting the number of sequence differences *m*_*i*_ in the haploid genotypes within the region, and diving by the region size *s*_*i*_ in base pairs: *θ*_*i*_ = *m*_*i*_/*s*_*i*_. The observed mismatch rate is then obtained by averaging all observations 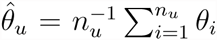, for *n*_*u*_ segments longer than *u* Morgans. We repeat this measurement for several thresholds *u*, obtaining a vector of observed mismatch rates 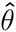. Due to the lack of detailed pedigree structures at deep time scales, the exact number of meiotic events separating two individuals that share IBD segments is generally unknown. Using the reconstructed demographic model, we therefore infer the posterior mean age *t*_*u*_ of pooled IBD segments longer than a known genetic length threshold *u*, using coalescent theory recently developed in [29, 30], the details of which are summarized in the Appendix. Finally, we regress the observed mismatch rates 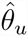 on twice the posterior mean age (in generations) to the MRCA of the IBD segments 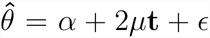. We refer to this regression as the tMRCA regression. Older segments will tend to harbor a larger number of sequence differences, due to the higher chance of mutation events occurring along the lineages connecting extant individuals to their most recent common ancestor. The slope *μ* of this regression will capture the rate at which mutations arise per unit of time. Note that we are neglecting the uncertainty on the measurement in the regressor *t*, i.e. the inferred age of the pooled IBD segments. As shown in simulations, however, this only results in negligible biases for the estimated slope coefficient due to the large number of pooled segments.

Assuming a genotyping error model for which false positive/negative genotype calls are independent of the average coalescent time of pairs of individuals at a locus, the intercept *α* of this regression is expected to capture the rate at which genotyping errors occur on the considered range of IBD segments.

### Controlling for gene conversion via MaAF-threshold regression

Non-crossover gene conversion events occur at a rate that is correlated to recombination, and have been observed to have frequency higher than recombination events [39]. In the coalescent process, gene conversion may be modeled as two consecutive recombination events that occur very close to each other [40], at an average distance of ∼300 base pairs [41, 42]. These events introduce the possibility that polymorphisms that are segregating in the population may be assimilated into haplotypes within IBD regions. These polymorphisms can create sequence differences between IBD individuals that are not due to newly arising mutations. Note that, whereas gene conversion events change the MRCA of the ∼300 bp converted segment, here we do not consider this to break an IBD block. Furthermore, because the number of gene conversion events is related to the number of meiotic events, short IBD regions will tend to exhibit more gene conversion-driven mismatches than longer, more recent IBD segments, therefore resulting in an upward bias when mutation rate is estimated via the slope of tMRCA regression. The mismatching variants observed on IBD segments, therefore, will be due to at least two distinct sources of heterozygosity. The first, *θ*_*p*_, which we call population heterozygosity in the remainder, represents the effect of gene conversion events which introduce standing genetic variation onto IBD blocks. The second source of heterozygosity is due to newly arising point mutations on IBD blocks, and will be referred to as *θ*_*μ*_. For IBD segments of a chosen length, we can express the total observed mismatch rate as *θ* = *θ*_*μ*_ + *θ*_*p*_. To estimate the mutation rate due to point mutations only, we need to exclude the effects of *θ*_*p*_ from our calculations. We make the following two observations:

1. The frequency of mutations that arise on long (e.g. *≥* 1 cM) IBD segments is typically low in the population (Figure S1), so that *θ*_*μ*_ is mostly due to rare variants.
2. If we divide the allele frequency spectrum into bins of equal width, we find an approximately uniform contribution to *θ*_*p*_ for each frequency. This implies that if we compute the frequency-bounded population heterozygosity *θ*_*p*,*f*_, using only variants of frequency at most *f*, we observe an approximately linear relationship between *θ*_*p,f*_ and *f* (Figure S2, see additional calculations in the Appendix).

Observation (1) implies that if we exclude high frequency variants when we compute *μ* using the proposed regression approach, the contribution of *θ*_*μ*_ to the observed mismatch rate on IBD segments will be largely unaffected. Furthermore, observation (2) suggests that if we estimate a frequency-bounded value of *μ*_*f*_ by ignoring variants of frequency higher than a threshold *f*, the contribution of population heterozygosity due to gene conversion events, *θ*_*p,f*_, will be decreased to an extent that is approximately linear in *f*. Assuming that the contribution of *θ*_*μ*_ to *μ*_*f*_ is unaffected for values of *f* in the range *F* = [*F*_*min*_, *F*_*max*_], we may therefore regress *μ*_*F*_ on *F*, and observe a linear relationship. We refer to this regression as the MaAF-threshold regression (Figure 2). The intercept of this regression will then reflect an estimate of *μ* without the confounding effects of *θ*_*p*_, while the contribution of *θ*_*μ*_ is left unchanged. We avoid computing values of *μ*_*F*_ corresponding to *F ∈* [0, *F*_*min*_), for a sufficiently large *F*_*min*_ (e.g. > 0.1), as this may result in removing variants that are due to new point mutation events on the IBD segments, which we use to estimate *μ*. Finally, note that we neglect the possibility that point mutations arising on IBD segments are removed via gene conversion, as this does not substantially affect the estimates.

### Estimating the gene conversion rate

The difference between the mutation rate computed without correcting for gene conversion events and the estimate obtained after removing the effects of gene conversion can be used to quantify the probability that a base pair within IBD segments is involved in a gene conversion event during meiosis. This difference, which we indicate as *μ*_*GC*_, represents the probability of observing a heterozygous site due to existing polymorphisms introduced via gene conversion in a single generation. This rate can be expressed as *μ*_*GC*_ = *p*(*GC*) × *p*(*θ*_*p*_*|GC*), i.e. the product of the probability of a base pair being involved in a gene conversion event, multiplied by the probability of assimilating a heterozygous site given the gene conversion occurs at the locus. The quantity *p*(*θ*_*p*_*|GC*) can be estimated using the genome-wide heterozygosity of the analyzed sample, and the value of *μ*_*GC*_ may be estimated using the previously described correction method. An estimate of *p*(*GC*) is therefore obtained as 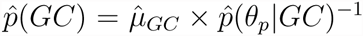, and a confidence interval is obtained having computed the standard error of these estimates via weighted block jackknife [43].

### Coalescent simulations

We used extensive coalescent simulation to evaluate the proposed methodology. To this end, we used a publicly available coalescent simulator, COSI2 [44], which allows simulating gene conversion events, and our implementation of a coalescent simulator, inspired by the existing GENOME algorithm [45], that enables simulations of a large number of samples and efficiently extract information on IBD segments. The algorithm proceeds backwards in time and, for each individual at generation *g*, samples a parent at the discrete time *g* + 1 in the past, occasionally resulting in coalescent events and sampling a new parent when a recombination event occurs. To speed up computation, the GENOME approach divides the simulated region into relatively large chunks that are not allowed to recombine, discretizing the recombination process and resulting in approximate LD structure at short genomic intervals. The version we developed enables substantial improvements of memory and run-time requirements, while circumventing the simplifying assumption of non-recombining LD blocks made by the original GENOME algorithm. Briefly, the speed-up over the original algorithm was obtained by sampling recombination breakpoints from an exponential distribution, and by only storing chromosomal regions and individuals that are relevant for the calculation of the ARG at each simulated generation. In addition to this, several improvements to data structures and other algorithmic details were applied. To evaluate our methodology, we further developed an extension of the program that allows efficiently extracting IBD segments from the ancestral recombination graph without requiring to test differences in shared common ancestors for each marginal tree in the ARG, as done in previous works [29, 38, 31]. We have released open-source software (ARGON) implementing the simulator (see Web Resources).

To asses the impact of demographic history on our estimates, we simulated three plausible demographic scenarios, in addition to the reconstructed GoNL demographic history. The simulated populations comprised an expanding population that experienced a severe founding event 30 generations before present, a population that undergoes severe exponential contraction (referred to as Ashkenazi and Maasai, respectively, due to resemblance with recently studied groups [29]), and an exponentially expanding population (referred to as Europeans, see Figure S3). We used two types of recombination maps to simulate non-uniform recombination rates along the genome. A map where the recombination rate alternates between 1cM/Mb and 2cM/Mb with intervals of 1 Mb, and one where one Mb every five has a six-fold increase in recombination rate over a baseline of 1cM/Mb (Figure S4). To asses the impact of genotyping errors on our methodology, we simulated three types of genotyping errors. We first simulated errors for which a previously unobserved variant is created (“de-novo” errors), or false-positive/negative calls on existing variants. To model frequency-dependent genotyping error rates, we used a beta distribution as a prior for sampling the frequency of planted genotyping errors [46]. For “de-novo” false positive errors, the frequency determines the number of individuals that are affected by an erroneous genotype call. For false-positive/negative genotyping errors, the sampled frequency corresponds to the frequency of the allele that is chosen to add/remove erroneous genotype calls. Three shape parameters were tested for the beta distribution: *α* = 0.01, *α* = 0.5, resulting in a strong preference for rare variants being erroneously called, and *α* = 1, resulting in a uniform distribution. In both cases the *β* parameter was set to 1 (see Figure S5). For all simulations, posterior mean estimates for the age of IBD segments were obtained using the coalescent distributions of the simulated models.

### GoNL data set

We analyzed sequence data from a recent study of 250 trio families from the Netherlands (the Genome of the Netherlands project [47], GoNL Release 4). The data set consists of 748 individuals that passed quality control. The samples were sequenced at an average of 13x using Illumina HiSeq 2000 technology, and variants were called using the GATK UnifiedGenotyper v1.6 software [48]. Trio phasing was performed using MVNcall [49], and additional quality control filters were applied as detailed in [47]. Indels (GoNL Release 5) were detected combining the output of several detection algorithms, as detailed in [47].

In addition to the quality control filters applied in the original analysis of the data, we further excluded regions that did not meet several quality criteria derived from the 1000 Genomes Project phase 1, as described in [50]. Specifically, we excluded from the analysis (1) low complexity regions; (2) markers that did not pass Hardy Weinberg equilibrium tests; (3) sites with excess coverage; (4) regions where common large insertions were detected; (5) regions not in the strict mask of the 1000 Genomes Project phase 1; (6) segmental duplications of the human genome.

Trio-phasing is expected to result in accurate estimation of haploid sequences in the GoNL data. Low frequency variants, in particular, are unlikely to result in doubly heterozygous parents, so that phasing of rare polymorphisms is generally trivial. Occasional phasing mistakes are however to be expected even in the presence of trios. For all analyses shown in the remainder, we have used a threshold of 1.0 for the MVNCall phasing and genotype calling posterior. This choice resulted in minimized estimated genotying error, with minimal effects on the estimated mutation rate (see Results, figures S6, S7, S8, and S9)

IBD segments and an inferred demographic model were obtained from the analysis described in [47]. For these analyses, only informative variants with very high trio-phasing and genotype calling quality were retained. For IBD detection, markers with frequency below 1% and with MVNCall [49] posterior less than 1.0 were excluded from IBD calculations, resulting in a total of ∼3. 500. 000 high quality variants. IBD detection was performed on haploid trio-phased individuals using GERMLINE [25] with parameters “-err hom 2 -err het 0 -bits 75 -haploid”, i.e. using windows of 75 markers, allowing a maximum of 2 mismatching sites per window to accommodate gene conversion and possible residual genotyping and trio-phasing errors, and retaining only segments of at least 1 cM for the demographic analysis. These parameters where chosen to be the most conservative parameters such that all transmitted haplotypes were detected intact in parent-child pairs along the genome. We further excluded from the analysis genomic regions outside 5 standard deviations from the mean genome-wide sharing, retaining 26 chromosomal regions, reported in [47], each longer than 45 cM, for a total of 2. 160 cM, and IBD density of 3.07 × 10^−3^ per site per pair.

Demographic inference was performed using the DoRIS software tool [29, 38]. The resulting demographic history is one of exponential expansion, with an ancestral population size of 11. 500 haploid individuals 150 generations in the past. Two periods of exponential expansion were inferred. The expansion rate between generations 150 and 10 was inferred to be 0.0146, followed by a strong expansion in the recent generations at rate 0.479 per generation. Due to the scarcity of extremely recent coalescent events, the magnitude of the latter expansion period is inferred with a high degree of uncertainty, however this was observed to not have appreciable effects on the analysis described in the remainder (see Results).

IBD detection is expected to be noisy at the boundaries of detected segments, so that non-IBD regions will be occasionally included in the estimated segments (and sometimes excluded). Because short IBD segments tend to harbor a larger fraction of miscalled nonIBD regions, which increase the observed mismatch rate, an upward bias in the tMRCA regression slope is expected. To cope with this, we have excluded 0.5 cM on either side of the IBD segments from the analysis of mutations and gene conversion rates, as we observed inflation due to noisy boundary estimation plateaus for values larger than this threshold (Figure S10).

Note that, due to violations of independence and homoscedasticity assumptions in the performed regressions, all reported standard errors were computed using weighted block jackknife [43], using the 26 independent chromosomal regions obtained as previously described.

### Enrichment of deleterious variation in IBD regions

We tested whether mutations arising between the present generation and the MRCA of IBD segments are enriched for deleterious variation. To this end, we ran the ANNOVAR software tool (version “2015Mar22” [51]) on the GoNL variants, and obtained numeric scores for the Polyphen 2 (“ljb23 pp2hvar” [52]) and Gerp++ (“gerp++gt2” [53]) annotations, restricting the analysis to scores > 2 for the latter. To test for enrichment, we compared the average score of genome-wide variants to the average score of variants found mismatching within IBD regions, treating all variants as independent and reporting Z-test p-values.

### Analysis of annotated genomic regions

Several sites along the genome were excluded from the analysis following the filtering criteria previously described. In addition, we analyzed mutation rates in specific regions described in several annotations (e.g. DNase I hypersensitive sites [54, 55, 56], histone modifications [57, 56, 58], constrained genes [4], and several others [59, 60, 61, 62, 61, 63, 64], see Table S1). It is sufficient to neglect regions that fall outside the genomic annotation at hand when computing the observed mismatch rate in the tMRCA regression. Annotations that are too small or too clustered in specific regions of the genome may result in downward biases of the estimated mutation rate, due to the “inspection paradox” of the Poisson process underlying the IBD sharing model [29] (Figure S11). This bias was computed and corrected using a permutation procedure (Table S1).

Sequence context is an important determinant of mutation rates, and trinucleotide context is often used to account for context-dependent mutation rate variation [65, 66]. When analyzing mutation rates within different genomic regions, we computed annotationspecific correction factors to account for the differences in mutation rates that are expected as a result of trinucleotide context variation, using the trinucleotide context-specific mutation-rate matrix of Kryukov [66] (details in Table S1).

We additionally derived mutation rates for different mutation categories: CpG/non-CpG and transition/transversions. First, we identified the ancestral allele for all GoNL variants. To do this, we downloaded the ancestral alignment used in the 1000 Genomes project ([67] see Web Resources). The ancestral allele for loci that were not present in this sequence (545, 279 out of 12, 181, 714) was set to the major allele found in the 1000 Genomes data set (*N* = 300, 503), or set to the allele found in the human reference genome hg19 (see Web Resources) if monomorphic in the 1000 Genomes data set (*N* = 244, 776). We then computed mutation rates using MaAF-threshold regression, excluding variants that did not match the analyzed mutation type (e.g. CpG transition), and scaled the resulting rate by the fraction of genome that may harbor the specific kind of mutation (e.g. CpG/non-CpG).

## Results

### Simulations

We evaluated the accuracy and robustness of the method via extensive coalescent simulation (see Materials and Methods). To assess the impact of demographic history on our estimates, we simulated several plausible demographic scenarios, and modeled genotyping errors using a beta distribution with different parameters, specifying error rate at different allele frequencies (Figure S5). We extracted ground truth IBD shared segments from the synthetic ancestral recombination graph, and simulated three types of errors, referred to as de-novo, false positive and false negative errors. De novo errors create erroneous variants in loci that are not truly polymorphic in the sequenced samples. False positive/negative errors affect existing variants, adding or removing derived alleles. To simulate frequency-dependent error rates, we sampled affected alleles using a beta distribution to select frequency of the derived allele for all kinds of errors. Inferred mutation rates under all three types of errors are displayed in Figure 3. We observed that tMRCA regression is robust to the presence of substantial levels of de-novo genotyping errors, consistent with the fact that IBD segments of different lengths are equally affected by the spurious sequence mismatches that result from errors of this kind. When false positive genotyping errors were simulated, we observed our approach to be robust to errors up to a rate of ∼10^−5^ per base pair. False negatives were tolerated up to a frequency of ∼10^−6^. Very large values of false positive/negative genotyping error rates resulted in a downward bias of the estimates, which is due to the fact that IBD segments of different lengths harbor a slightly different spectrum of mismatching sites, and are therefore not equally likely to be affected by spurious genotype calls (see Figure S1). Similar results were observed for several kinds of genotyping error distribution, demographic model, and recombination map, although the approach proved more robust for error distributions that are less concentrated on very rare variants (figures S3, S4, S5, and S12). The intercept of the tMRCA was observed to reflect genotyping error, with average values between 1 and 2 times the simulated error rate (Figure S13), depending on the type of error and the parameters of the distribution used to select the frequency of affected alleles.

**Figure 3:**
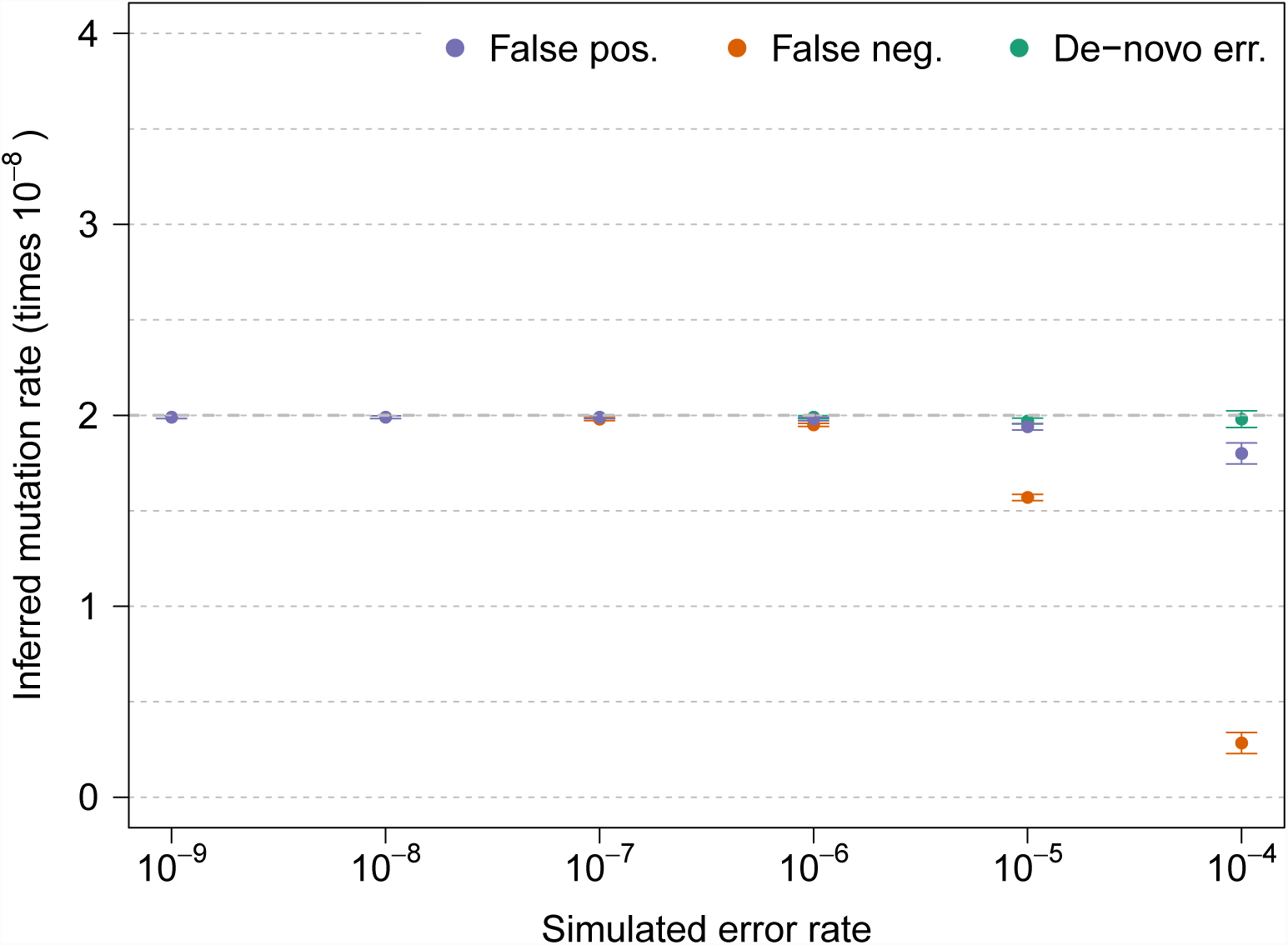
Inferred mutation rates under several values of simulated genotyping error rate, for three types of genotyping errors. The simulated true underlying mutation rate was *μ* = 2 × 10^−8^. All simulations involved a single chromosome of 250 cM for 200 haploid individuals from a GoNL-like population using *Beta*(*α* = 0.5, *β* = 1) as a prior for allele frequency of erroneous variants. True IBD segments were extracted from the simulated ancestral recombination graph. Additional simulation results are shown in Figure S12.

To compare the power of the proposed method to the power of trio-based mutation rate inference, we simulated data at various sample sizes using the GoNL demographic model. Due to the quadratic increase in IBD sharing pairs as sample size increases, the proposed method results in smaller standard errors than the trio-based approach, except at very small sample sizes (Figure 4). However, for demographic models that result in substantial IBD sharing due to a small recent effective population size, higher sample size did not substantially decrease the standard error (Figure S14). This is due to the fact that as new samples are added, early coalescent events result in overlapping ancestral lineages across pairs of individuals, so that limited new information is obtained from increasing the sample size.

**Figure 4:**
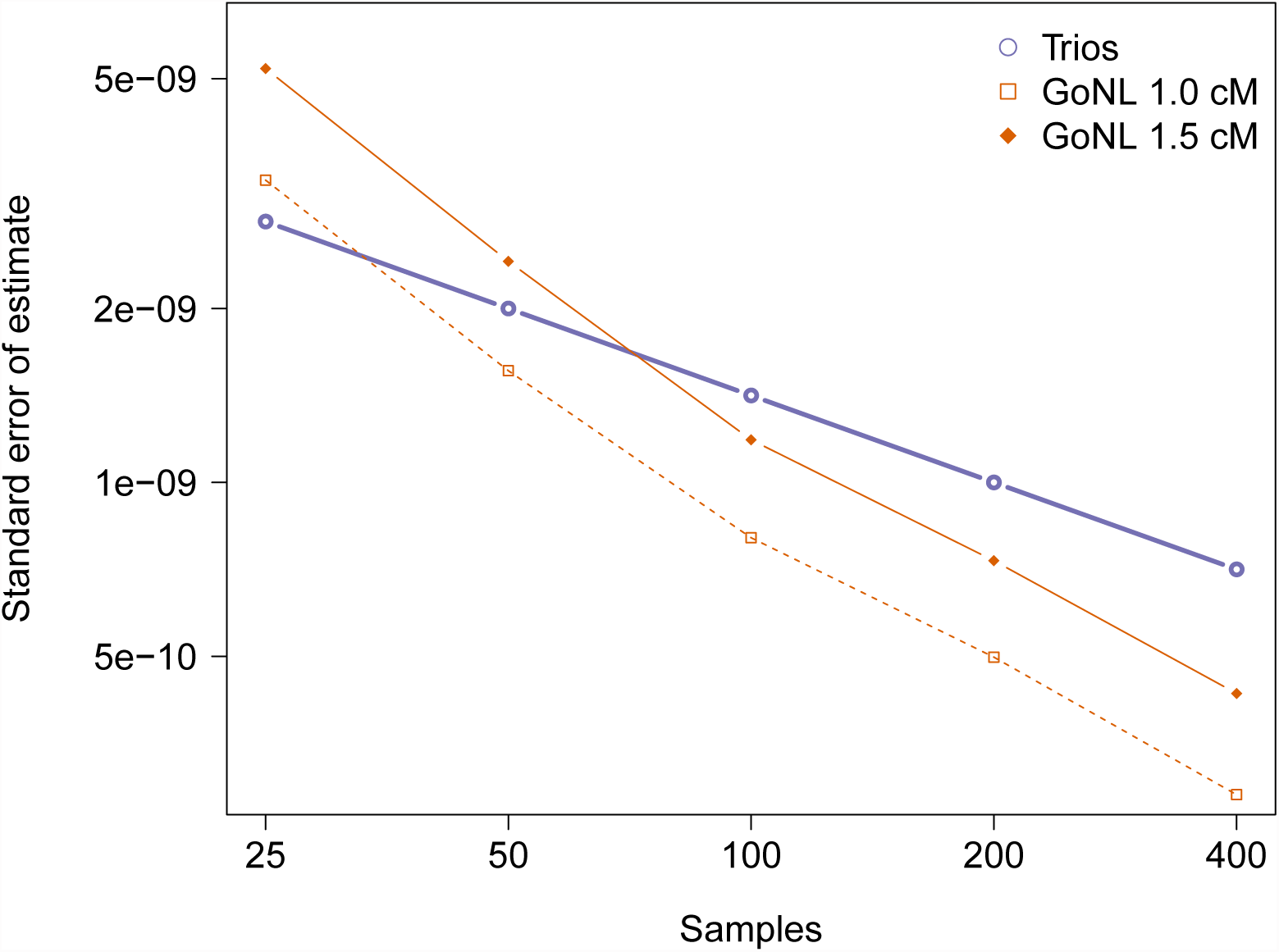
Comparison of the estimate standard error for trios and tMRCA under different demographic models and minimum IBD segment length cutoffs. We report the estimated standard deviation from the analysis of several simulations of a single 100 Mb chromosome. For illustrative purposes, we show results of analyses using IBD length cutoffs of 1.0 and 1.5 cM. Analysis of the GoNL data was performed using a length cutoff of 1.6 cM.

We finally tested the MaAF-threshold regression approach to correct biases introduced by non-crossover gene conversion events and estimate the probability that a base pair is involved in gene conversion. We simulated realistic mutation and gene conversion rates and used GERMLINE to detect IBD sharing after subsampling synthetic SNPs in order to match the allele frequencies observed in the GoNL data. We observed good performance of the MaAF-threshold-regression in recovering the simulated mutation rate value (Figure 5), and a small downward bias when recovering the gene conversion rate using the GERMLINE IBD discovery parameters used in the real data analysis (Figure S15).

**Figure 5:**
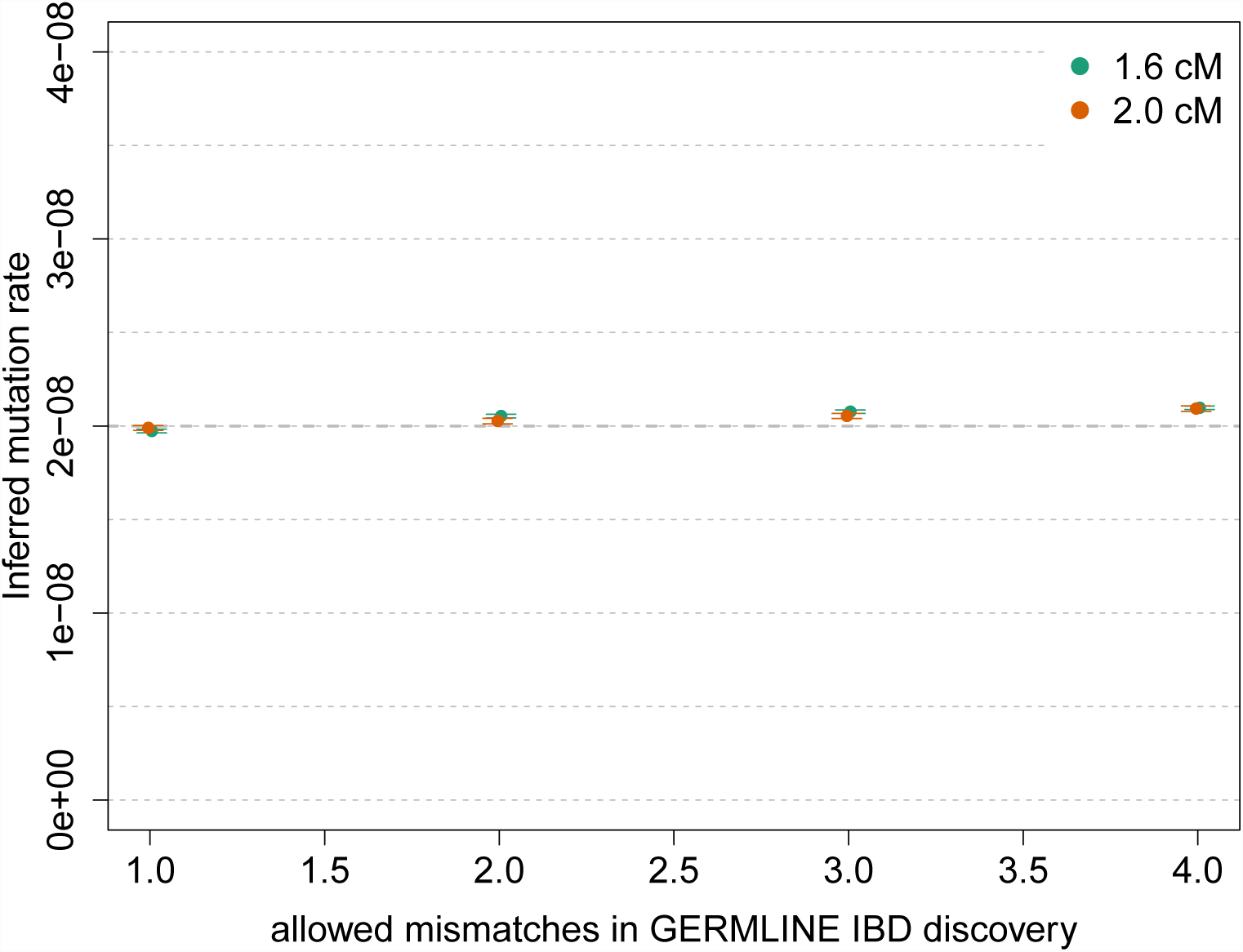
Inference of gene conversion-corrected mutation rate in simulated data. We simulated a chromosome of 50 cM for 250 diploid samples, using *μ* = 2 × 10^−8^ for the mutation rate and a probability of 6 × 10^−6^ for a basepair to be involved in a non-crossover gene conversion event. We matched the allele frequency spectrum of the simulated samples to the spectrum found in real data for IBD detection with GERMLINE. We used several values of the GERMLINE allowed mismatching sites (“-het”) to asses the impact of this parameter in the results. Negligible biases are observed for the recovered mutation rate.

### Average genome-wide mutation rate and gene conversion rate in the GoNL data set

We analyzed 498 founders that passed quality control in 250 trio families sequenced within the Genome of the Netherlands (GoNL) project (see Materials and Methods). Due to the trio design of the GoNL study, the average ∼13*×* sequencing depth is effectively doubled to ∼26*×* for the transmitted haplotypes in the 498 analyzed founders. 248 trios and 2 duos passed sequencing quality control. In the remainder, we report results for the analysis of transmitted haplotypes only.

We estimated a mutation rate of 2.08 ± 0.06 × 10^−8^ (Figure 6) before correcting for gene conversion events. For all analyses of mutation and gene conversion rates, we report results for a minimum IBD segment length of 1.6 cM, discarding 0.5 cM on either edge of the segments, and ignoring variants with a trio-phasing and genotyping posterior value less than 1.0. Choosing more conservative values for the minimum length and edge exclusion cutoffs resulted in compatible estimates (Figures S10, S16, and S17). As expected, including variants with lower trio-phasing and genotyping posterior resulted in higher estimates of genotyping error, but negligible effects were observed on the estimates of mutation rate (figures S6, S7, S8, and S9). The tMRCA regression intercept, which reflects genotyping and phasing error rate (see Materials and Methods), was estimated to be 2.21 ± 0.09 × 10^−6^, within a range that is not expected to result in biases in the tMRCA regression slope based on simulations (figures 3, S6, S9, S12, and S13).

**Figure 6:**
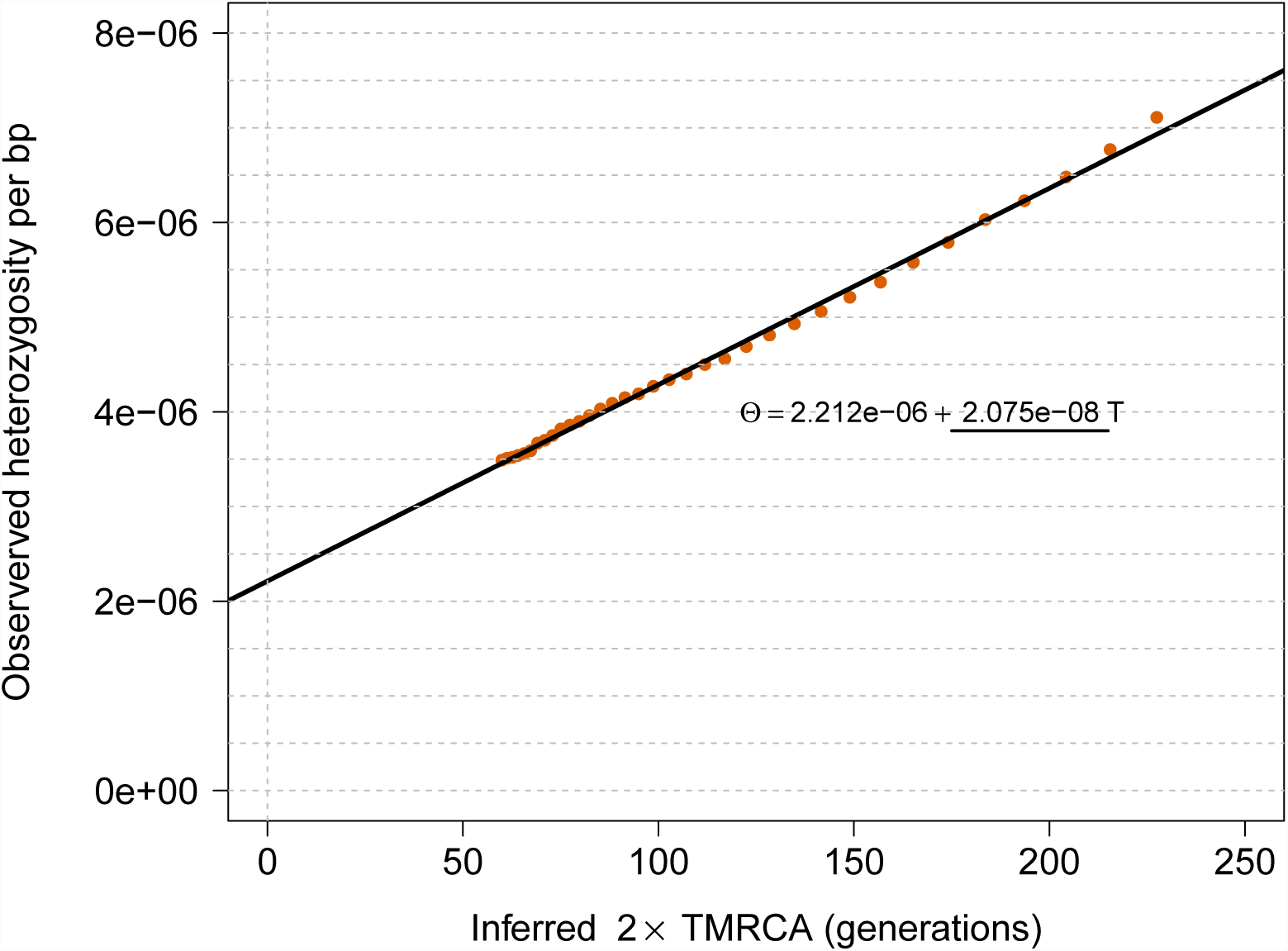
tMRCA regression for segments of length *≥* 1.6 cM in the GoNL data set. The obtained slope is used to estimate mutation rate per generation per base pair, before the effects of gene conversion are accounted for.

We then performed MaAF-threshold regression to correct for gene conversion events (see Materials and Methods, Figure 7). Using this approach, we estimated a genomewide average mutation rate of 1.66 ± 0.04 × 10^−8^ per base, per generation. The difference between the corrected and uncorrected estimates, 4.18 ± 0.48 × 10^−9^, reflects the chance of a heterozygous base pair entering IBD segments during meiosis, as a result of non crossover gene conversion. This quantity can be used to estimate the chance that a base pair is involved in a gene-conversion tract (see Materials and Methods). Heterozygozity in the GoNL data is observed to be ∼6.98 × 10^−4^, consistent with a diploid long-term effective population size in humans of approximately 10, 500 individuals. Using this quantity, a base pair is estimated to be involved in a gene conversion event at a rate of 5.99 ± 0.69 × 10^−6^ per meiotic event. This rate is in good agreement with a recently published estimate of 5.9 ± 0.71 × 10^−6^ [39]. We further computed estimates of the rate of mutations at CpG and non-CpG sites (Table S3). The obtained estimates for CpG sites were higher than in previous reports based on trio analysis, consistent with a higher genome-wide rate (see Table 2 in [20]).

**Figure 7:**
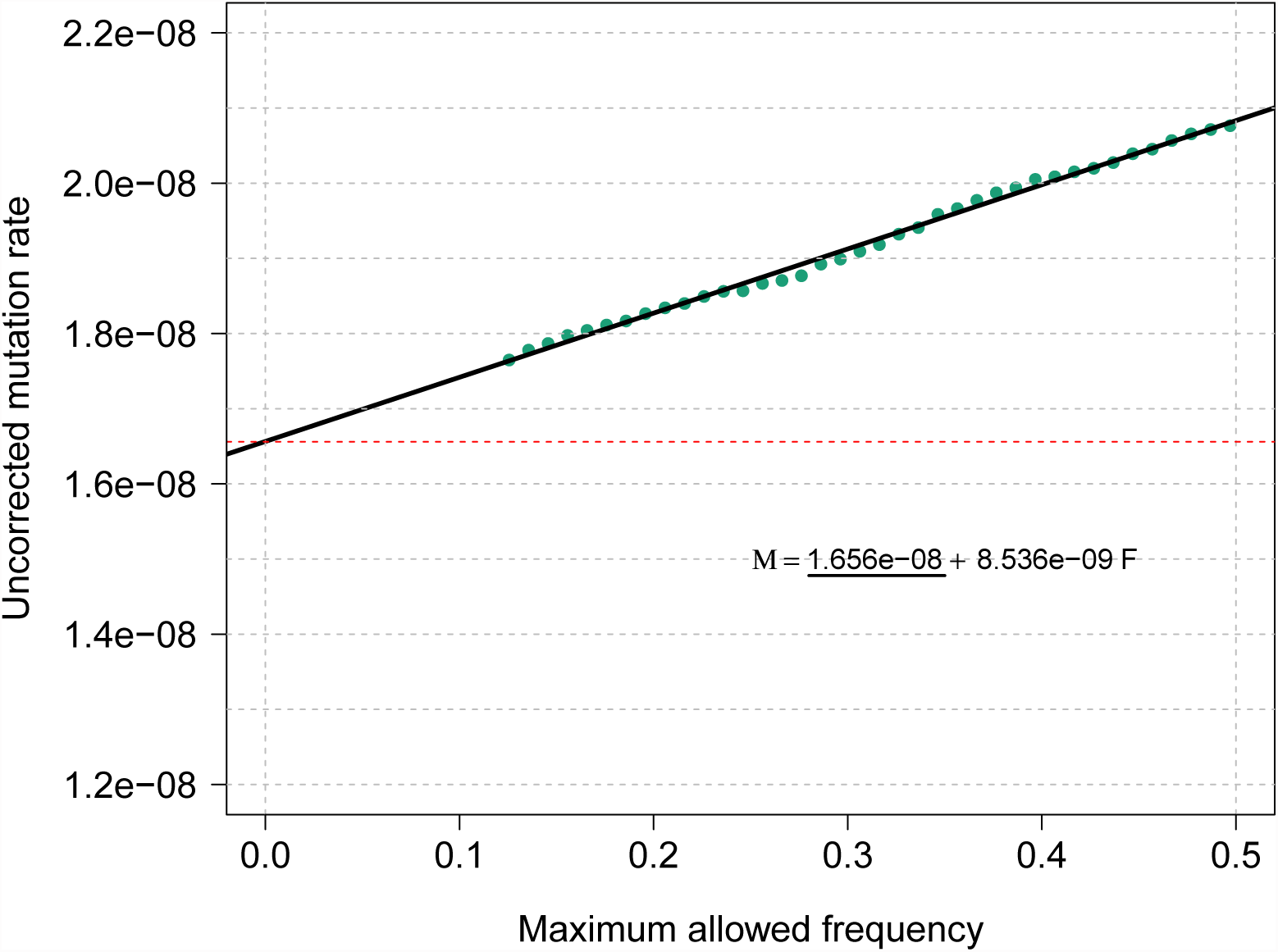
MaAF-threshold regression for segments of length *≥* 1.6 cM in the GoNL data set. We compute mutation rates for several allowed maximum allele frequency thresholds between 0.125 and 0.5 (green dots), and regress the observed heterozygosity on the maximum allele frequency. The intercept of the resulting linear model reflects the corrected mutation rate estimate.

Because our analysis relies on a reconstructed demographic model, which is used to determine IBD segment age, we performed sensitivity analysis to asses the impact of potential inaccuracies of the reconstructed model on our results (Table S2). The demographic history of the analyzed Dutch samples comprises two consecutive periods of exponential expansion (Figure S3). When a genome-wide average mutation rate was inferred for a demographic model with ancestral population size perturbed by 10%, we observed a ∼1.8% difference in the inferred average mutation rate. We observed very limited effects on the mutation rate estimate when perturbing the present-day population size, which is inferred with uncertainty due to the scarcity of very recent coalescent events (Table S2).

### Average genome-wide indel rate

Besides measuring the rate of germline point mutation events, the proposed approach allows quantifying additional evolutionary parameters associated with the transmission of IBD chromosomal regions. We applied the same procedure used for inferring mutation rates to infer the rate of < 20bp indels, which we estimated to be 1.26 ± 0.06 × 10^−9^. This rate is higher than a recent estimate of 0.68 × 10^−9^ reported in [68], but compatible with a second recent estimate of 1.5 ± 0.18 × 10^−9^ [21], both obtained via observation of de-novo events in trios. We additionally used our method to estimate the gene conversion rate based on indels, obtaining a rate of 9.02 ± 2.91 × 10^−6^ per meiotic event, compatible with the rate obtained from point mutations.

### Recombination does not strongly impact mutation rate

We used our approach to analyze annotation-specific mutation rates (see Materials and Methods). We looked for association between recombination rates and mutation rates, a relationship that has been previously detected and attributed to mutagenic properties of recombination [33]. Indeed, we found our tMRCA regression estimates of mutation rate to be strongly associated with recombination rate (*β* = 0.38 ± 0.04 mut/rec, *p* = 5.27 × 10^−6^, R-squared = 0.9, Figure 8). As previously mentioned, however, increased sequence mismatch rate at loci that undergo frequent recombination may be a result of polymorphic variants introduced by gene conversion events, which may increase the slope of the tMRCA regression. Consistently, after controlling for gene conversion, we observed no significant association between recombination rate and mutation rate (*β* = −0.04±0.03, *p* = 0.17), suggesting the lack of observable mutagenic effects associated with recombination hotspots (figures 8 and S18). A recent study reached similar conclusions [7]. Repeating the same analysis with indels, we detect no significant association between indel rate and recombination rate, for both tMRCA regression slope and MaAF-threshold regression intercept (*β* = −0.003 ± 0.003, *p* = 0.37 after gene conversion correction).

**Figure 8:**
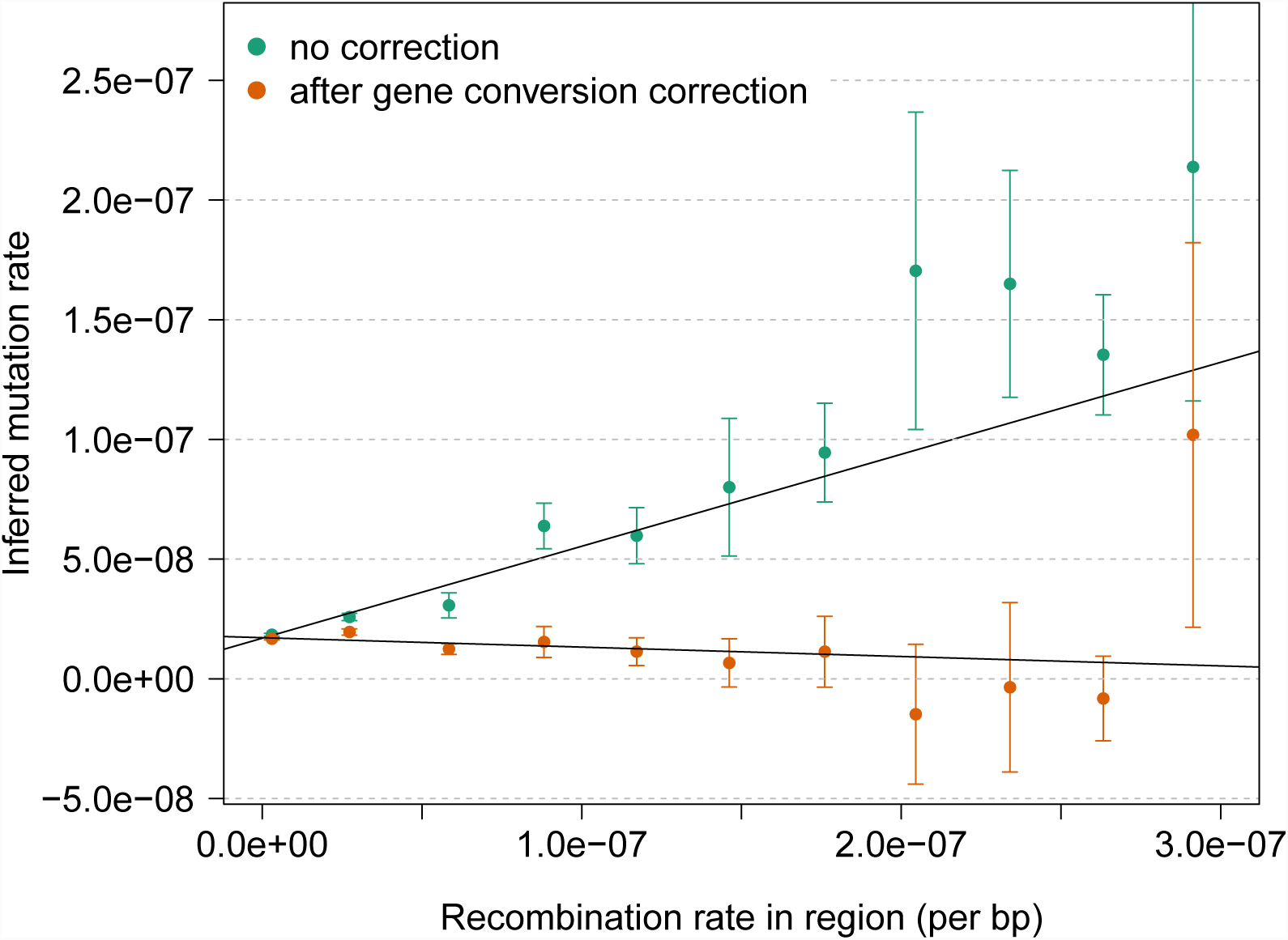
Association between recombination rate and mutation rate. We annotated the genome based on uniform bins of recombination rate, and estimated mutation rates for each obtained annotation. We observed a strong association between mutation and recombination rate before correcting for the occurrence of gene converion events. After applying the correction we detect no significant association, which suggests the linear relationship observed for the uncorrected estimates is induced by gene conversion (See Figure S18).

### Effects of background selection

Natural selection affecting new mutations may reduce genomic variation, leading to downward bias in our mutation rate estimates. Because our analysis is limited to mutation events that occurred in the past ∼100 generations, due to the length of IBD segments, we expect the effects of natural selection on our genome-wide average mutation rate estimate to be small. Genomic regions with functional or regulatory roles, however, may be under selective pressures that may result in measurable impact even at these short time scales.

To estimate the impact of selective pressures on our estimates, we divided the genome based on the B statistic proposed in [5]. The B statistic measures the impact of background selection on a genomic region by estimating the ratio between local effective population size and the effective population size expected under neutrality, so that small values of the B statistic correspond to higher selective pressures (See [5], page 11 for details on the computation of the B statistic). Similarly, a local reduction in effective population size impacts the spectrum of IBD shared segments, which are expected to be longer on average, as a result of early coalescent events in populations of smaller effective size [69, 37]. Indeed, we observed a strong correspondence between small values of the B statistic and the average length of IBD segments (*p* = 8.43 × 10^−7^, Figure 9). The effect is, as expected, such that smaller values of the B statistic correspond to longer average IBD shared segments, due to reduced local effective population size. This effect is remarkably strong up to the measured genome-wide average value of the B statistic. We observed longer average IBD segments for large values of the B statistic, a result that may be explained by biases in either of the two measures, or by the fact that additional evolutionary forces, such as selection acting on standing genetic variation [69], are being captured by IBD segment lengths. When we measured the impact of different values of the B statistic on our estimates of mutation rate, however, we found the effect to not be significant (*β* = 2.17 ± 1.55 × 10^−9^ mutations per generation, per unit of B statistic, *p* = 0.19, Figure S19). The average genome-wide value for the B statistic was estimated to be 0.78 in the analyzed regions, suggesting a moderate amount of background selection is acting at the average analyzed locus (a value of 1 reflects absence of background selection). If we were to correct the estimated average genome-wide mutation rate to account for this, we would obtain an updated average mutation rate of 1.7 ± 0.05 × 10^−8^, which is however not significantly different from the estimate obtained without accounting for background selection.

**Figure 9:**
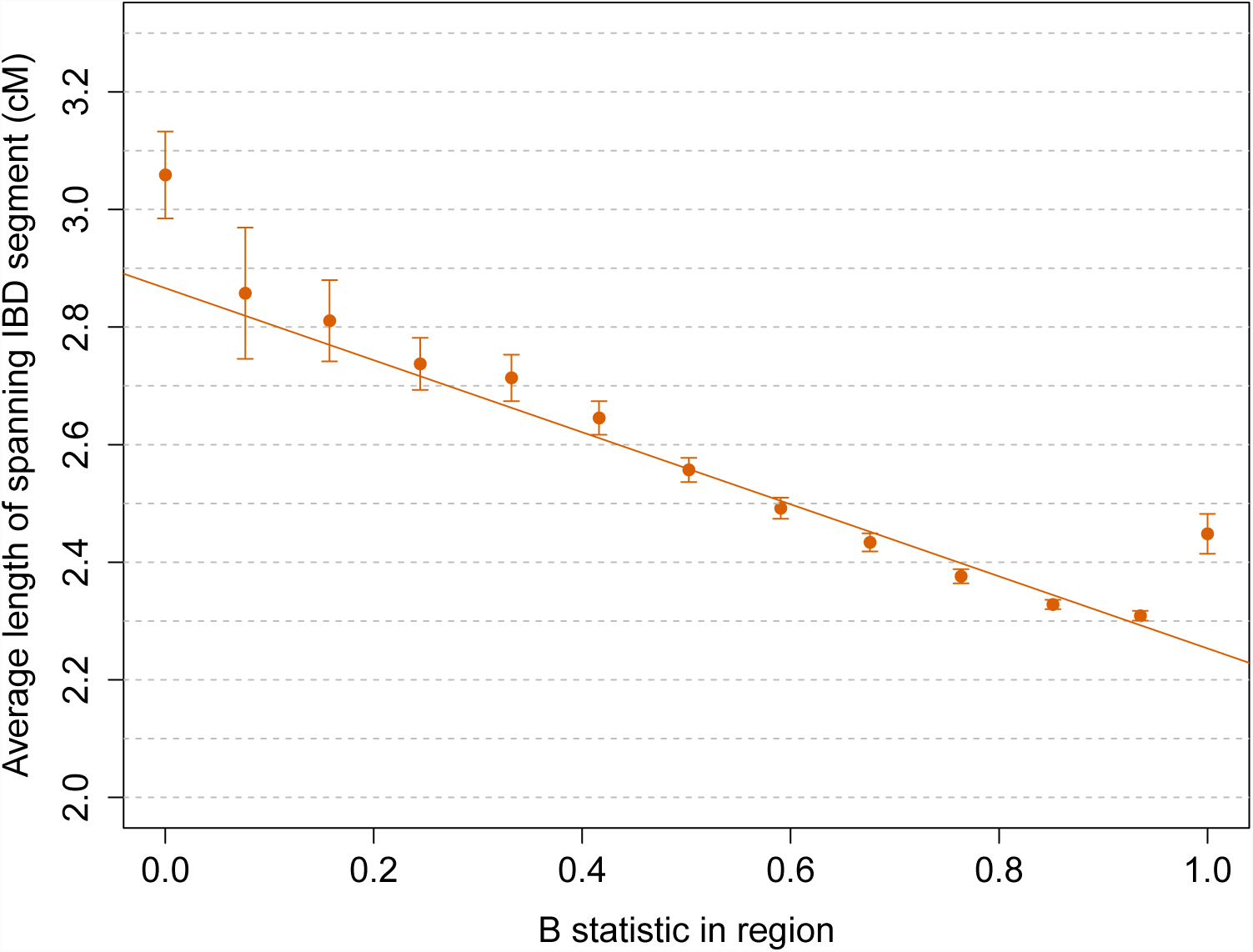
Relationship between region-specific values of the B statistic and the average length of IBD segments > 1.6 cM spanning the regions. Equally-spaced bins of the B statistic were used. Reduced local effective population size has similar effects on the B statistic and the length of IBD haplotypes, which are longer in regions of strong background selection due to earlier average coalescent times between pairs of individuals.

### Sequence differences in IBD segments are enriched for deleterious variation

Mutation events occurring within the analyzed IBD regions are expected to have arisen within the past ∼100 generations, and are therefore on average substantially younger than variants randomly sampled along the genome. Several recent studies have outlined the recent origin of a large fraction of functionally relevant variants [70, 71, 72, 73, 74]. We therefore tested whether the presence of recent mutations on IBD segments resulted in an enrichment of deleterious variants compared to the average genome-wide locus, contrasting average scores obtained using Polyphen 2 [52] and Gerp++ [53] annotations (see Materials and Methods). Of the analyzed GoNL variants, 54, 960 were annotated using Polyphen 2, and 948, 782 were annotated using Gerp++. Of these, 1, 843 and 27, 900 were found mismatching on IBD segments of 1 cM or longer, respectively. When average scores were compared, we found that mismatching sites within IBD regions were strongly enriched for higher scores in both annotations (Table 1, Polyphen2 Z-test *p* = 2.8 × 10^−5^; Gerp++ Z-test *p* = 9.03 × 10^−10^). We further found a marginal association between Ployphen 2 scores and the B statistic of background selection (*β* = −0.074 ± 0.025, *p* = 0.014, R-squared = 0.39) and a strong association between Gerp++ scores and regional B statistics (*β* = −0.734 ± 0.068, *p* = 3.55 × 10^−7^, R-squared = 0.91, Figure S20), which is expected due to both measures relying on metrics related to sequence conservation.

**Table 1:**
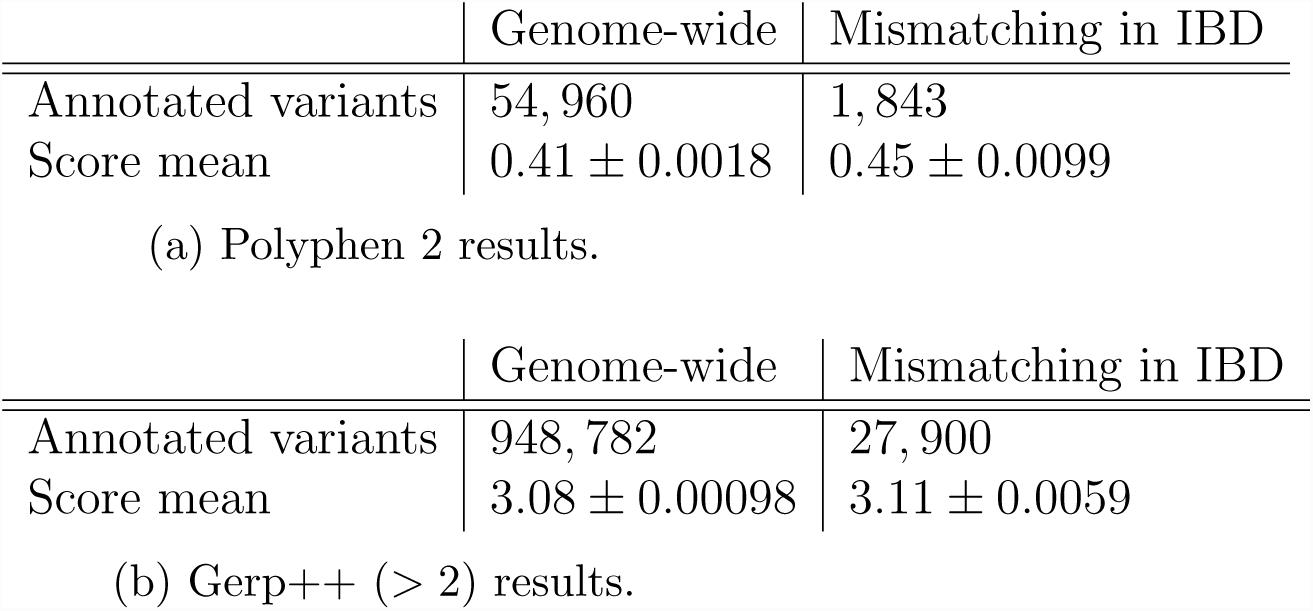
Analysys of Polyphen 2 and Gerp++ annotated variants: genomewide vs. mismatching within IBD segments.

We finally tested for enrichment/depletion of the mutation rate in several genomic annotations that have recently been extracted from several studies (e.g. DNase I hypersensitive sites [54, 55, 56], histone modifications [57, 56, 58], constrained genes [4], and several others [59, 60, 61, 62, 61, 63, 64]). Annotation-specific mutation rates after correcting for the effects of trinucleotide context (see Materials and Methods) are reported in Table S1. None of the annotations were significantly enriched or depleted for mutation rates after controlling for trinucleotide context and multiple hypothesis testing. A recent paper [75], found that cell-specific chromatin features are a strong determinant of cancer mutations. Our estimated mutation rate in DNase I hypersensitive regions of 1.66 ± 0.05 × 10^−8^, on the other hand, suggests that the germline mutation rate is not substantially different from the genome-wide average in these regions, in line with recent analyses [12].

## Discussion

Because mutation events play a key role in the shaping of heritable variation, accurate characterization of this evolutionary parameter is a central task of data-driven analyses of genomic data. In this paper, we proposed a new method to infer mutation rates using distant relatives in a large group of purportedly unrelated individuals, using long chromosomal regions co-inherited identical-by-descent by present day individuals. The proposed method relies on recently developed coalescent calculations to infer population size history at very recent time scales, which is in turn used to reconstruct the distribution of ages for IBD segments detected in the population. This procedure has several advantages compared to existing other methods. First, by regressing observed sequence mismatch rates on IBD segment age, the estimation is robust to substantial amounts of genotyping error, which is an important confounding factor for many recent estimators of mutation rates based on trio data. Second, by modeling the relationship between population heterozygosity and allele frequency, it is possible to remove the effects of non-crossover gene conversion, while at the same time estimating its rate. In addition, this approach can be used to estimate the rate of other kinds of mutation events, such as insertions and deletions.

We inferred a genome-wide average point mutation rate of 1.66 ± 0.04 × 10^−8^ per base per generation, or 5.71 ± 0.14 × 10^−10^ per base per year, having assumed a generation length of 29 years [76]. Recent family-based estimates range within 1.0 − 1.2 × 10^−8^ per base per generation [10, 13, 14], being in most cases significantly lower than the estimate we report here. These methods have the advantage of relying on direct observation of de-novo mutation events, with minimal modeling assumptions, but are affected by the need to rely on strict filtering criteria to deal with false positive/negative genotype calls, which may explain the discrepancy with our results. Phylogenetic methods, on the other hand, fall within 2.0 − 2.5 × 10^−8^ [15, 16]. These estimates rely on several underlying modeling assumptions, which provide a possible explanation for the higher inferred rates, although some have suggested the possibility that these analyses may be capturing the results of evolutionary changes of the mutation rate, or the effects of a varying length of generation times [10, 13, 77]. The estimate of 0.4 − 0.6 × 10^−9^ per year computed in [22] using ancient DNA is slightly lower than our result, but the reported confidence intervals are compatible. Similarly, the rate of 1.4 − 2.3 × 10^−8^ per generation reported in [78], and computed based on point-mutations nearby microsatellites is compatible with our estimate.

A contemporary study [23], which is related in spirit to ours, used simulation-based calibration of the decay of heterozygosity along the genome to infer an average genomewide mutation rate of 1.65 ± 0.1 × 10^−8^. This closely matches our estimated value. The authors discuss several implications of this mutation rate on our ability to reconcile demographic events inferred using DNA analysis and fossil records, which apply to our analysis as well. Because conversion between sequence divergence and phylogenetic split times across different primate species relies on a per-year mutation rate estimate, different values of this rate have a direct impact on our ability to reconstruct the timing of these events [10, 13]. In general, lower values of the mutation rate (e.g. the pedigree-based “slow” mutation rate) result in older estimates of demographic events, while larger values (e.g. the “fast” phylogenetic rates) result in earlier demographic events. Our inferred “intermediate” mutation rate value of ∼5.71×10^−10^ per year, consistent with that of [23], implies intermediate scalings for the timing of these events which are generally compatible with fossil records. Assuming no significant effects of generation time and no changes in mutation rates, our estimate implies the split between humans and chimps occurred ∼6.6 million years in the past, and the split between humans and orangutan about ∼16 m.y. in the past. When our estimate of mutation rate is used to interpret recently reported split times across human populations [11], we find dates that are compatible with what has been reconstructed using methods other than DNA-based reconstruction. The split of African and non-African populations is estimated to have occurred 46 − 61 thousand years in the past, while a split time of 15 thousand years is inferred for the separation of East Asians and Native American populations. These estimates are lower than those obtained assuming a “slow” mutation rate, but do not contradict current fossil evidence.

In addition to estimating the rate of point mutations, we report a gene conversion rate of 5.99 ± 0.69 × 10^−6^, in close agreement with a recent report [39], and find that recombination is not associated with mutation rates, supporting recent findings [7]. A recent sperm-typing study further dissected the relationship between mutation, recombination and gene conversion, finding evidence for higher mutational load in regions of high recombination, which is however contrasted by repairing mechanisms associated with gene conversion [79]. These lead to a higher prevalence of GC alleles compared to AT alleles. Overall, these effects may be counteracting each other in a way that results in minimal differences in the total number of observed mutations in recombination-rich regions, while affecting sequence composition. Interestingly, a recent study reports that recombination rate affects the distribution of putatively deleterious variants along the genome, but found no evidence for a role of biased gene conversion in this observation [80].

Finally, our method was applied to estimate the rate of short (< 20 bp) indels, which have not thus far been extensively characterized. We inferred a rate of 1.26 ± 0.06 × 10^−9^, compatible with two previous estimates of 1.5 ± 0.18 × 10^−9^ [21], and 1.06 ± 0.1 × 10^−9^ [81], but higher than the estimate of 0.68 × 10^−9^ reported in [68]. While these analyses are likely affected by difficulties related to detection of short indels, collectively they suggest that insertion and deletion mutation events occur at a significantly lower rate compared to single point mutations.

In addition to analyzing genome-wide average rates, we looked for enrichment or depletion of mutation rates in a number of genomic annotations that were recently derived from several studies. Although we cannot exclude significant deviations from genome-wide averages, we found no evidence for changes in overall mutation rates for the analyzed regions. Notably, we observed that while the distribution of IBD shared haplotypes closely reflects the effects of background selection along the genome, a negligible effect is observed on our estimated mutation rates, suggesting that estimating mutation rates using mutation events under the effects of ∼100 generations of natural selection does not significantly bias local mutation rate estimates in European populations. Consistent with the idea that mutations on IBD segments are recent and under the effects of selective forces [70, 71, 72, 73, 74], we found a strong enrichment for deleterious variants within IBD regions.

Our method provides a new way of studying mutation and gene conversion events in large samples of unrelated individuals, being robust to substantial amounts of genotyping error, which constitutes a significant confounder in trio-based analyses. The main limitation of our approach, however, is the need to rely on two fundamental components that are potential sources of bias, namely detection of shared IBD segments and the need to infer the recent demographic history for the analyzed population. Our analysis of mutation rates in the GoNL dataset relies on IBD detection and demographic inference performed in a previous study [47], but it is possible that additional sources of uncertainty in these two components affect our results. Our conservative exclusion of substantial portions of IBD segments, together with our sensitivity analysis for changes in the demographic model, however, suggest that these biases, if present, should not be substantial.

Several potential directions for improvement of the proposed methodology and analysis may be outlined. First, additional developments of the coalescent calculations used in this work may remove the requirement of estimating a demographic model for the analyzed samples, as described in [82]. These methods need to be adapted in order to deal with genotyping error and to deconvolute the contribution of mutation and gene conversion to observed sequence mismatches in IBD regions. Second, it may be possible to devise improved approaches for dealing with heteroscedasticity and the dependence across observations in the tMRCA and MaAF-threshold regressions, which do not result in biases, but may reduce the efficiency of our estimators. In addition, the MaAF-threshold regression currently assumes an underlying model of constant effective population size, an approximation that only has small effects on our estimates (Figure S2), but may be removed via additional modeling. Third, as shown in our simulation-based evaluation of the method, the relationship between allele frequency and genotyping error rates limits the robustness of our method to very large amounts of noise in the analyzed sequences. Alternative genotype calling strategies may be employed to reduce these effects, e.g. genotype calling approaches that do not rely on whether variants are observed polymorphic in other sequenced individuals. Finally, by applying the proposed tMRCA regression, it may be possible to analyze multi-generation pedigrees while controlling for substantial genotyping error. Detection of very long IBD segments is in fact trivial when closely related individuals are analyzed, and the age of these segments is fully determined by their known genealogical relationship. In addition, the proposed methods can be applied to increasingly large datasets that are currently being produced, which may be reliably phased to enable accurate IBD detection even in the absence of sequenced family members [24].

## Appendix

### The age of IBD segments

If a pair of chromosomes find a common ancestor at time *t* generations before present, the probability that a single site is spanned by an IBD segment of length *l* at least *u* Morgans can be expressed as

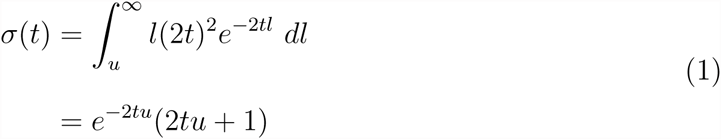

The distribution *l*(2*t*)^2^*e*^−2*tl*^ represents the sum of two exponential random variables with parameter 2*t*, which is the rate at which a recombination occurs on either side of the chosen site. Note that this assumes an IBD segment is delimited by the occurrence of recombination events, which is equivalent to assuming an underlying sequentially Markovian coalescent (SMC) model. For very short IBD segments (e.g. < 0.3 cM) and in populations that experience substantial and long-lasting isolation (e.g. *N*_*e*_ < 1, 000), the slightly more complex SMC’ model [83] provides more accurate calculations [84, 85, 11, 86]. This is however unnecessary given the demographic history and length ranges here considered. It follows from the linearity of the expectation operator that the expected fraction *f* (*t*) of genome shared IBD for a pair of individuals whose ancestral lineages coalesce at time *t* can be obtained from the probability that a single site is spanned by an IBD segment of length at least *u* Morgans, which we write *f* (*t*) = *σ*(*t*). The expected length of an IBD segment transmitted from a common ancestor living at time *t* is therefore

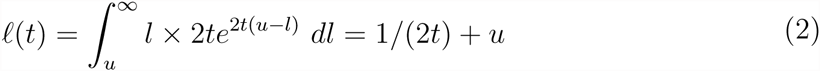

To obtain the expected number of IBD segments obtained if the lineages of two individuals coalesce at time *t*, we therefore divide the expected total amount of genome shared IBD by the expected length of an IBD segment co-inherited from an ancestor living at time *t*. This yields

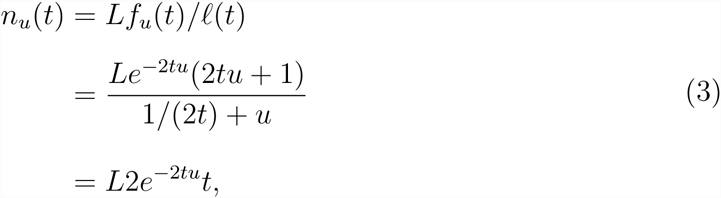

where *L* is the size, in Morgans, of the considered genomic region. To obtain the expected number of IBD segments longer than *u* Morgans for the average pair of individuals in the population, we marginalize over the distribution of pairwise coalescence times, *c*(*t*), which depends on the demographic history,

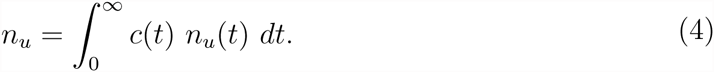

This quantity has a closed form expression if we assume that the population size becomes constant at an arbitrarily remote point in time, and can be used to obtain the posterior age distribution of IBD segment ages,

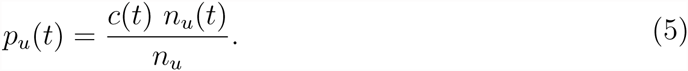

### Contribution of individual variants to heterozygosity

For a sample of *K* homologous sequences from a population, the heterozygosity per site can be estimated by computing

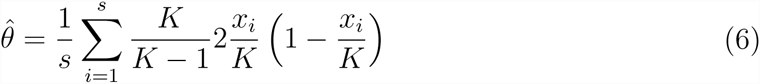

[87], where *s* is the number of sites in each sequence, *x*_*i*_ is the number of samples carrying a derived allele at site *i*, and *K/*(*K -* 1) is a bias-correction factor. Defining *M* (*x*) as the total number of sites in the sample for which exactly *x* sequences carry a derived allele, we can rewrite this equation as a sum over *x*:

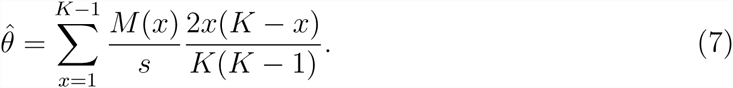

The term *M* (*x*)*/s* is the proportion of sites at which *x* of the *K* sequences carry a derived allele and the term

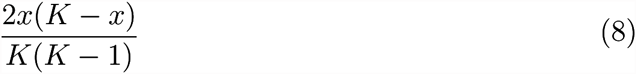

is the probability of discovering such a polymorphic site when just two sequences are sampled without replacement from the *K* sequences. Note that this probability is the same for sites with *x* copies of a derived allele as it is for sites with *K - x* copies. Thus, we may also write

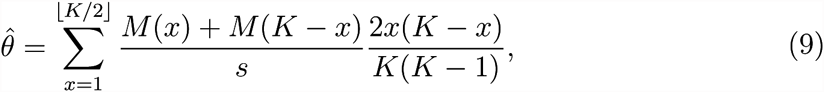

where *[K/*2] is the largest integer that is less than or equal to *K/*2 and *x* is now the count of the minor allele.

This allows us to consider the average contribution of different kinds of polymorphic sites to overall heterozygosity. Under the constant population size, neutral model of [88],

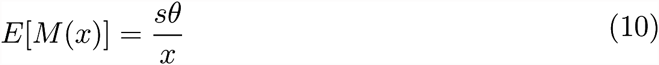

[89, 90] in which *θ* = 4*N μ* is the diploid population-scaled mutation rate per site, or the expected per-site heterozygosity of the population. Using (10) together with (9) and simplifying gives

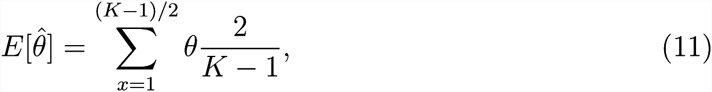

in which we have assumed that *K* is odd for simplicity. The sum in (11) evaluates to *θ*, as expected for an unbiased estimator.

Equation (11) shows that on average the different kinds of polymorphic sites, categorized by minor allele frequency, contribute uniformly to heterozygosity, as noted previously by [91]. Another way of stating this is that polymorphisms discovered by screening in samples of size two will be uniformly distributed among minor-allele frequency classes. This depends on the population size being constant over time, and is also not true for derived allele-frequency classes. Figure S2 shows that contributions to heterozygosity are close to uniform for the GoNL site frequency spectrum even though the population size has not been constant.

## Supplemental data

Supplemental Data include 20 figures and 3 tables.

## Acknowledgments

We express our gratitude to Nick Patterson, Priya Moorjani, Mark Lipson and Amy Williams for useful scientific discussions and comments on an early draft, and to Ilya Shlyakhter for support with the COSI2 simulator. This research was funded by NIH grant R01 MH101244.

## Web Resources

The developed coalescent simulator (ARGON) will be made available at https://github.com/pierpal/ARGON.

The tool to infer mutation and gene conversion rates (IBDMUT) will be made available at <u>https://github.com/pierpal/IBDMUT</u>.

The 1000 Genomes ancestral alignments were downloaded from ftp://ftp.1000genomes.ebi.ac.uk/vol1/ftp/phase1/analysis_results/supporting/ancestral_alignments/.

The human reference sequence (h19) was downloaded from http://hgdownload.cse.ucsc.edu/goldenPath/hg19/chromosomes/.

## Consortia

The members of the Genome of the Netherlands Consortium are Laurent C Francioli, Androniki Menelaou, Sara L Pulit, Freerk van Dijk, Pier Francesco Palamara, Clara C Elbers, Pieter B T Neerincx, Kai Ye, Victor Guryev, Wigard P Kloosterman, Patrick Deelen, Abdel Abdellaoui, Elisabeth M van Leeuwen, Mannis van Oven, Martijn Vermaat, Mingkun Li, Jeroen F J Laros, Lennart C Karssen, Alexandros Kanterakis, Najaf Amin, Jouke Jan Hottenga, Eric-Wubbo Lameijer, Mathijs Kattenberg, Martijn Dijkstra, Heorhiy Byelas, Jessica van Setten, Barbera D C van Schaik, Jan Bot, Isac J Nijman, Ivo Renkens, Tobias Marschall, Alexander Schnhuth, Jayne Y Hehir-Kwa, Robert E Hand-saker, Paz Polak, Mashaal Sohail, Dana Vuzman, Fereydoun Hormozdiari, David van Enckevort, Hailiang Mei, Vyacheslav Koval, Matthijs H Moed, K Joeri van der Velde, Fernando Rivadeneira, Karol Estrada, Carolina Medina-Gomez, Aaron Isaacs, Steven A McCarroll, Marian Beekman, Anton J M de Craen, H Eka D Suchiman, Albert Hofman, Ben Oostra, Andr G Uitterlinden, Gonneke Willemsen, LifeLines Cohort Study, Mathieu Platteel, Jan H Veldink, Leonard H van den Berg, Steven J Pitts, Shobha Potluri, Purnima Sundar, David R Cox, Shamil R Sunyaev, Johan T den Dunnen, Mark Stoneking, Peter de Knijff, Manfred Kayser, Qibin Li, Yingrui Li, Yuanping Du, Ruoyan Chen, Hongzhi Cao, Ning Li, Sujie Cao, Jun Wang, Jasper A Bovenberg, Itsik Pe’er, P Eline Slagboom, Cornelia M van Duijn, Dorret I Boomsma, Gert-Jan B van Ommen, Paul I W de Bakker, Morris A Swertz & Cisca Wijmenga

